# A Multi-Tissue Comparison and Molecular Characterization of Canine Organoids

**DOI:** 10.1101/2022.07.15.500059

**Authors:** Christopher Zdyrski, Vojtech Gabriel, Oscar Ospina, Hannah Wickham, Dipak K. Sahoo, Kimberly Dao, Leeann S. Aguilar Meza, Abigail Ralston, Leila Bedos, William Bastian, Sydney Honold, Pablo Piñeyro, Eugene F. Douglass, Jonathan P. Mochel, Karin Allenspach

**Affiliations:** SMART Pharmacology, Department of Biomedical Sciences, Iowa State University, Ames, IA, USA; SMART Pharmacology, Precision One Health Initiative, University of Georgia, Athens, GA, USA; 3D Health Solutions Inc., Ames, IA, USA; Department of Biostatistics and Bioinformatics, Moffitt Cancer Center, Tampa, FL, USA; Department of Veterinary Clinical Sciences, Iowa State University, Ames, IA, USA; Pharmaceutical & Biomedical Sciences, Institute of Bioinformatics, University of Georgia, Athens, GA; Veterinary Diagnostic Laboratory, Iowa State University, Ames, IA, USA

**Author notes:** **Corresponding authors:** Christopher Zdyrski Ph.D.; K. Allenspach DVM, Ph.D.; Jonathan P. Mochel DVM, Ph.D. Authors contributed equally to the manuscript (Christopher Zdyrski, Vojtech Gabriel). co-senior authors (Jonathan P. Mochel, Karin Allenspach). **DECLARATION OF INTERESTS** K. Allenspach is a co-founder of LifEngine Animal Health and 3D Health Solutions. She serves as a consultant for Ceva Animal Health, Bioiberica, LifeDiagnostics, Antech Diagnostics, Deerland Probiotics, and Mars. J.P. Mochel is a co-founder of LifEngine Animal Health and 3D Health Solutions. Dr. Mochel serves as a consultant for Ceva Animal Health, Boehringer Ingelheim, Dechra Ltd. and Ethos Animal Health. C. Zdyrski is the Director of Research and Product Development at 3D Health Solutions. K. Dao was an employee of 3D Health Solutions. A. Ralston was an employee of 3D Health Solutions. Other authors do not have any conflict of interest to declare. Multiple patents related to this work have been filed through the Iowa State University Research Foundation.

**Keywords:** canine, dog, organoids, reverse translational medicine, stem cell, endometrium, lung, pancreas, bladder, liver

## Abstract

Organoids are 3-dimensional (3D) stem cell-derived cell culture lines that offer a variety of technical advantages compared to traditional 2-dimensional (2D) cell cultures. Although murine models have proved useful in biomedical research, rodent models often fail to adequately mimic human physiology and disease progression, resulting in poor preclinical prediction of therapeutic drug efficacy and toxicity. With the advent of organoid technology, many of these challenges can be overcome. Previously, the use of canine organoids in drug testing and disease modeling was limited to organoids originating from the intestine, liver, kidney, lung, and urinary bladder. Here, we report the cultivation, maintenance, and molecular characterization of two novel adult-stem cell-derived canine organoid cell lines, including the endometrium and pancreas, in addition to previously reported bladder, lung, and liver organoids from two genetically related canines. Five tissues and organoid lines from each donor were characterized using bulk RNA-seq, allowing for a unique, multi-organ comparison between these two individuals and identification of specific cell types such as glandular epithelial cells in endometrial organoids.

## INTRODUCTION

Numerous *in vitro* models are used as preclinical biological and pharmacological research tools (Hickman et al., 2014). The most prevalent *in vivo* models for biomedical research include *Drosophila melanogaster* and *Caenorhabditis elegans*, both of which are widely used for genetic research, *Danio rerio*, a model used primarily for mutagenesis screening, and numerous mammalian models for which advanced genetic tools are available (Kim et al., 2020b). Mouse models are extensively used in biomedical research due to their cost-effectiveness, fast-growing nature, and availability of genetic mutants (Kim et al., 2020b). Differences in diet, living environment, circadian rhythm, and the short lifespan are among the issues limiting the translational relevance of rodent models (Perlman, 2016). Although the murine model has proven effective in a variety of biological research areas, rodents frequently fail to adequately mimic human physiology and disease progression, hence compromising their predictive performance in preclinical pharmaceutical research (Gordon et al., 2009; Wang et al., 2018a). Approximately ninety percent of experimental drugs fail to make the transition from discovery to successful clinical trials (Mochel et al., 2018; Van Norman, 2016; Shalev, 2006). Drug development is successful in about 5% of those ultimately approved, and average costs including post-approval R&D in the U.S. exceed $2 billion and it takes 10-15 years to reach clinical trials per drug (Brancato et al., 2020; DiMasi et al., 2016). The use of 3D organoids in the screening stage of drug discovery could drastically reduce the use of live animals for drug development (Mollaki, 2021). Ultimately, additional research is warranted to identify alternative *in vitro* models that can more accurately replicate human physiology and reduce animal use. Conventional pharmacology research involves using 2D cell culture and animal testing prior to human clinical trials (Brancato et al., 2020).

Organoids are 3D self-organized, miniature, and simplified versions of organs *in vitro*. Adult stem cells can self-renew, differentiate into multiple cell types, and are genomically stable over multiple passages (Fatehullah et al., 2016; Huch and Koo, 2015; Huch et al., 2015). Unlike traditional 2D cell lines, organoids grow in a 3D extracellular matrix, allowing for the recreation of more realistic tissue architecture and physiological responses (Hynds and Giangreco, 2013). Recently, donor-to-donor variability in human ileum- and colon-derived organoids has been investigated; however, more research is needed across other biomedical models, including dogs (Mohammadi et al., 2021). Furthermore, organoids can be used in both basic and applied biomedical research, including the study of genetic disorders, cancers, and infectious diseases (Kim et al., 2020b; Nantasanti et al., 2015; Usui et al., 2017; Zhou et al., 2018). Organoids can be a valuable tool for personalized medicine where patient-specific organoids can be grown and incubated with drug candidates to predict effectiveness prior to patient treatment (Bartfeld and Clevers, 2017; Kaushik et al., 2018). Organoids may become useful for regenerative medicine; for example, hepatic organoids can be transplanted to a patient and transdifferentiated to various hepatocellular regional identities *in vivo* (Sampaziotis et al., 2021). Organoid cell lines drastically reduce the number of animals needed for drug testing as they can be expanded indefinitely in culture and cryopreserved for future use. Organoid technology has made it possible to undertake pharmacological research and testing in a manner that is more responsible from an ethical standpoint, while also simplifying the process of genetic manipulation (Artegiani et al., 2020; Takeda et al., 2019). While human organoids are a valuable research tool in the biomedical field, they come with limitations. Public concern plays a key role in tissue sampling from human patients (Lehmann et al., 2019). Ethical concerns for the use of human-derived organoids include chimeric research and genetic editing of organoids derived from patients (Munsie et al., 2017). Although a growing number of studies have characterized cell populations using transcriptomic data across human organs, there is a lack of similar studies in non-human models (Jones et al., 2022).

The reverse translational paradigm, in which data from human clinical research might aid in the development of veterinary therapeutics and vice versa, is garnering a growing amount of interest (Schneider et al., 2018). Dogs share similar lifestyles and diets with their owners due to the close relationship between dogs and humans, often including a sedentary lifestyle and an increased risk of developing obesity (Chandler et al., 2017). The longer lifespan of dogs over that of mice predisposes dogs to develop analogous chronic diseases to humans, including diabetes mellitus (Adin and Gilor, 2017), ocular diseases (Sebbag et al., 2018, 2019), inflammatory bowel disease (Chandra et al., 2019; Jergens and Simpson, 2012; Kopper et al., 2021), congestive heart failure (Mochel and Danhof, 2015; Mochel et al., 2019; Silva and Emter, 2020), cancers (Knapp et al., 2020), and cognitive dysfunction (Ozawa et al., 2019), among others (Wang et al., 2018a). Therefore, in addition to being used as a large animal model for preclinical drug safety assessment, dogs are also emerging as a translatable model for demonstrating proof-of-concept efficacy studies, particularly in the field of oncology (LeBlanc et al., 2016; Schaefer et al., 2016). While dogs excel as a model in many applications compared to rodents, they come with their own challenges in the form of expensive housing and ethical concerns about using live dogs in research. Dogs are recognized as companion animals in western countries, and there are ongoing worldwide initiatives to limit their use in research through the 3Rs (*Reduce*, *Replace*, *Refine*) principles (Hasiwa, 2011; Russell and Burch, 1959). A potential solution that provides more access to the canine model while decreasing reliance on live animal use lies in organoid technology.

Organoids still represent a relatively novel technology and lack formal standardization in isolation, maintenance, and downstream applications. We previously demonstrated the ability to culture canine intestinal organoids from healthy and diseased tissues and demonstrated the translational potential of these organoids for human medicine (Chandra et al., 2019). Our research group has been working to standardize the culture and maintenance of organoid cell lines to maximize the reproducibility of our findings across different experimental sites (Gabriel et al., 2022a). This investment into the standardization of protocols includes downstream applications such as the use of a permeable support system for canine organoids in drug testing and discovery (Gabriel et al., 2022b), use of organoids in viral testing (García-Rodríguez et al., 2020), and regenerative medicine (Sampaziotis et al., 2021).

Currently, there is a lack of canine organoid models compared to other major biomedical species to accurately depict and study various diseases, drugs, and biological phenomena. This report describes the successful cultivation of two novel canine organoid lines, including the endometrium and pancreas, to the best of the authors’ knowledge, there has not been any peer-reviewed description of these organoids, in addition to previously described bladder (Elbadawy et al., 2022), lung (Shiota (Sato) et al., 2023), and liver (Nantasanti et al., 2015) organoids from two related dogs. By comparing five tissue-specific organoid lines obtained from two genetically related donors (B816 and B818), this study aims to acquire insight into gene expression in different organoids and their corresponding tissues. Preliminary analyses using RNA sequencing, immunohistochemistry, and immunofluorescence were used to characterize the relationships between individual organoid cell lines and their parent tissue. In addition to characterization, we compared related individuals across organs in this preliminary analysis, thus preparing these organoids to next be utilized and tested in a variety of biomedical applications and functional assays.

## RESULTS

### Organoid Expansion

Organoid cell lines were successfully established from five organs, including the uterus, lung, pancreas, urinary bladder, and liver, two of which are novel, originating from two female canine individuals. All tissues were isolated on the same day, minced, and embedded in Matrigel. Organoids were cultivated simultaneously, and growth progression, passage number, and media are reported in **Supplemental file 1.** Samples were passaged between two and four times to remove excess tissue fragments before being harvested for characterization and freezing (**Figure 1-figure supplement 1**). Wells with any remaining tissue were excluded from any characterization to ensure reliable results. The same media was used for all organoid lines and the media composition is listed in **Supplemental file 2** and consists of growth factors which encourage the growth of multiple tissues. Several of the canine organoids displayed a variety of distinct morphological phenotypes characterized via light microscopy and hematoxylin & eosin (H&E) staining (**Figure 1**).

**Figure 1.**
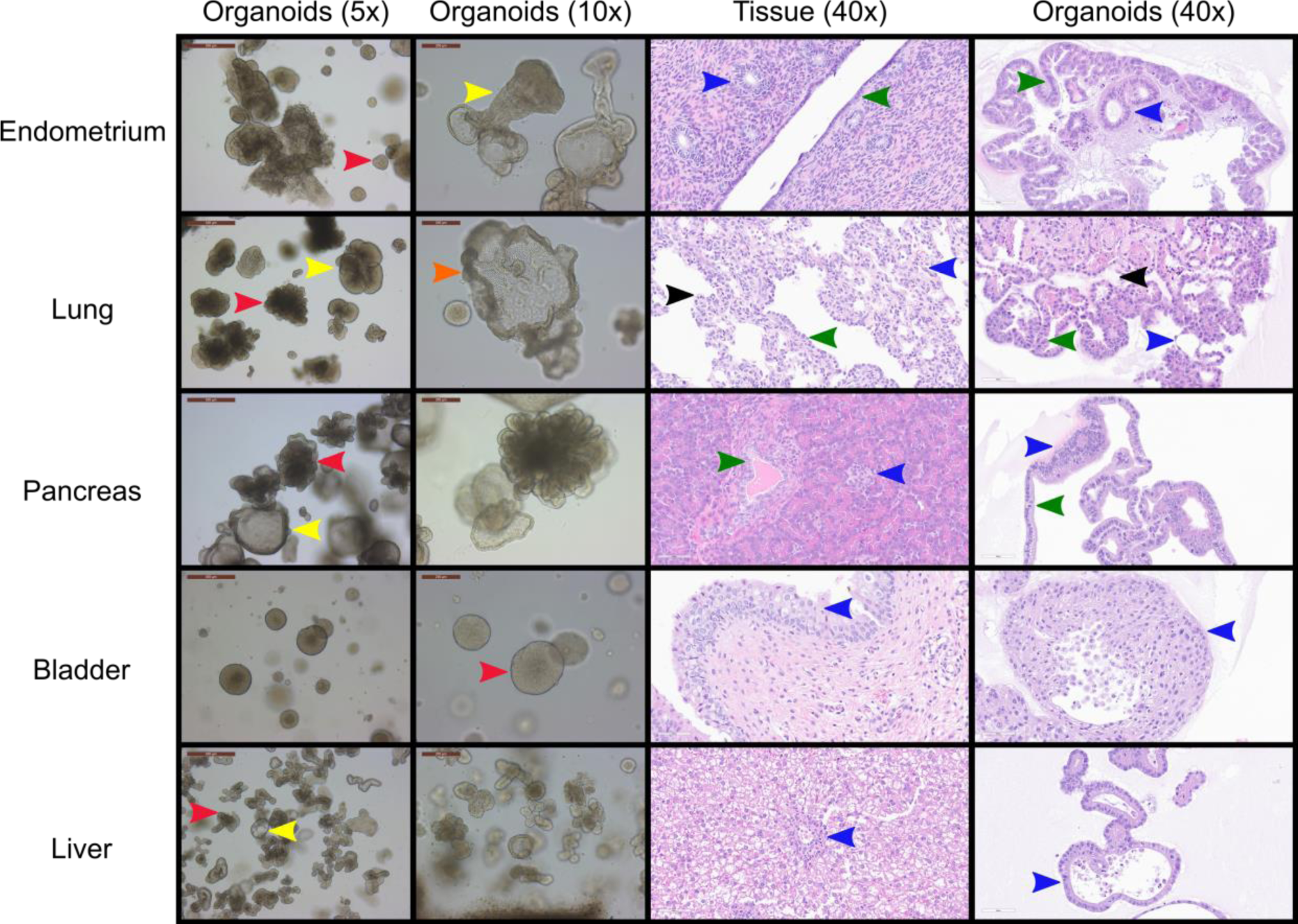
Morphological and histological characterization of canine organoid lines derived from a single donor. Bright field and hematoxylin & eosin (H&E) images and approximate proportions of the cultures for the five organoid cell lines derived from the uterus (red=80%, yellow=20%), lung (red=60%, yellow=35%, orange=5%), pancreas (red=50%, yellow=50%), bladder (red=100%), and liver (red=80%, yellow=20%) of canine individual B816. Red, yellow, and orange arrows indicate distinct morphologies in each organoid line while blue, green, and black arrows indicate similar histological areas of the organoids and tissues. Structures identified in tissues with histological similarities in organoids were found in endometrium (blue=glandular epithelial cells, green=endometrial epithelial cells), lung (blue=alveolar type-1 cells, green=bronchial epithelial cells, black=alveolar type-2 cells), pancreas (blue=endocrine cells, green=intercalated ducts), bladder (blue=transitional epithelium), and liver (blue=cholangiocyte) organoids. Images of organoid cultures were captured using a Leica Dmi1 microscope. Scale bars are provided at 5X (500 μm), 10X (200 μm), and 40X (60 μm) magnifications.

Immunohistochemistry (IHC) for Pan cytokeratin (PanCK) was used to confirm the epithelial origin of the canine organoids, while tissues were positive only in epithelial regions (**Figure 2**). To better confirm and distinguish that the organoids were not remaining tissue, smooth muscle actin (SMA) was stained (**Figure 2**) and vimentin (VIM) was used to identified mesenchymal cells in the tissues and organoids (**Figure 2**).

**Figure 2.**
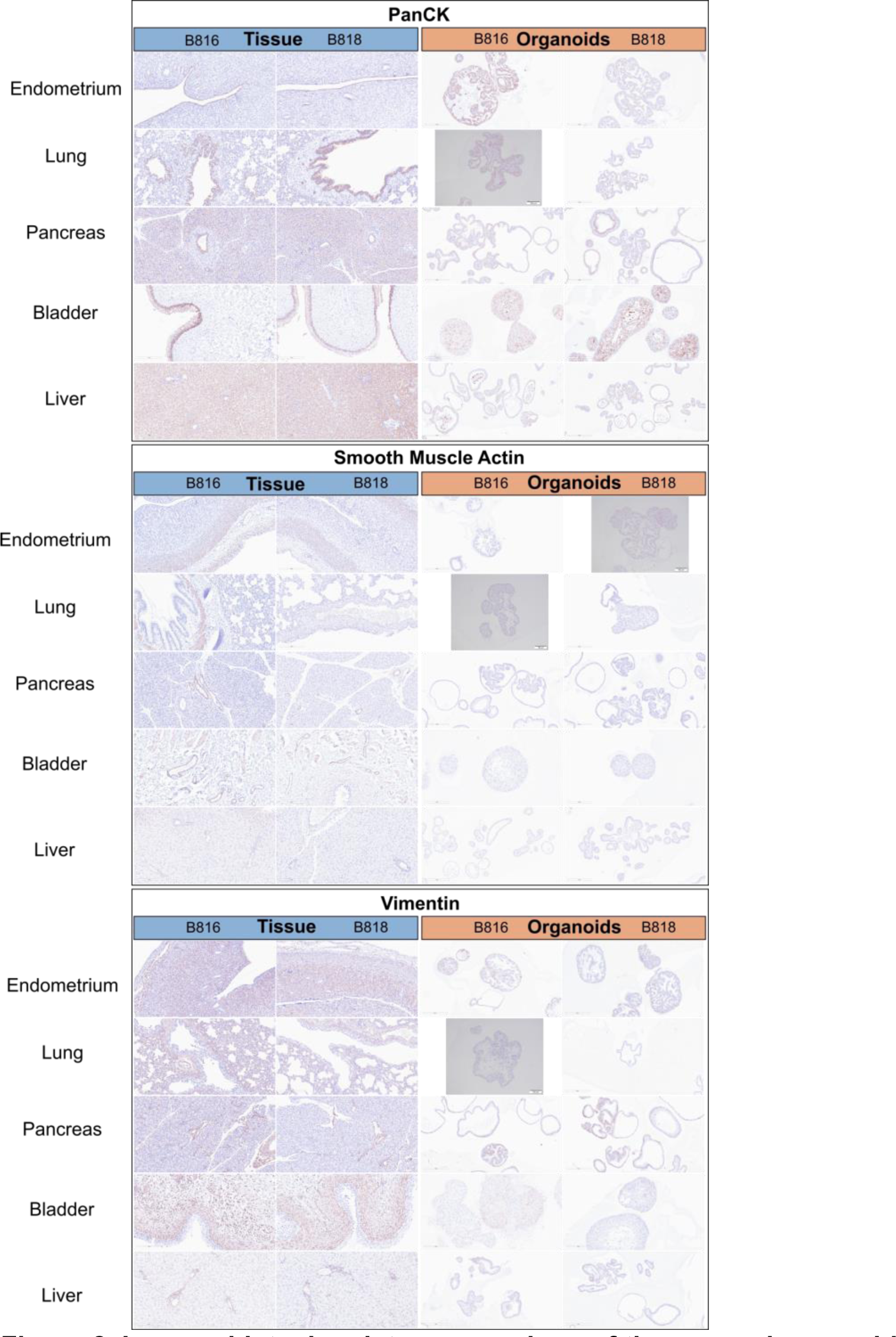
Immunohistochemistry comparison of tissues and organoids. Immunohistochemistry staining of organoids and tissues for both donors, B816 and B818. Scale bars for each image are displayed in μm. Antibodies for pan cytokeratin, smooth muscle actin, and vimentin were used. Images of negative control slides are provided in **Figure 2-figure supplement 1**.

### Morphological and histological characterization of canine organoid cell lines

#### Uterus

A subset of endometrial organoids formed a tubular structure appreciated on brightfield microscopy during culture (**Figure 1**). H&E staining suggested that the culture consists of endometrial epithelial cells and glandular epithelial cells (**Figure 1**). RNA-seq data indicated that SRY-box transcription factor 17 (SOX17) was upregulated in endometrium organoids (**Figure 3B**). This protein is expressed in the human endometrium, specifically in the luminal and glandular epithelium (Kinnear et al., 2019). SOX17 is important for endometrial glandular development and function in mice (Turco et al., 2017), and was also expressed in organoids derived from human menstrual flow, consistent with their function in endometrial gland development (Cindrova-Davies et al., 2021). Uroplakin Ib (UPK1B) was upregulated in endometrium organoids compared to uterus tissues (**Figure 3A**). The top endometrium-specific genes for organoids included Actin gamma 1 (ACTG1), Clusterin (CLU), and Prothymosin alpha (PTMA), whereas uterine tissues showed high expression of multiple ribosomal proteins (**Figure 3D**). Regarding endometrium-specific genes, intra-organoid comparison revealed 1,039 unique genes (**Figure 4A**), uterine intra-tissue comparisons revealed 2,930 unique genes (**Figure 4B**), and 10,487 genes were expressed in both organoids and tissues (**Figure 4C**). For immunofluorescence (IF), Vimentin (VIM) was positive in both tissues and organoids while Paired Box 8 (PAX8) identified endometrial glands in tissues (**Figure 5**).

**Figure 3.**
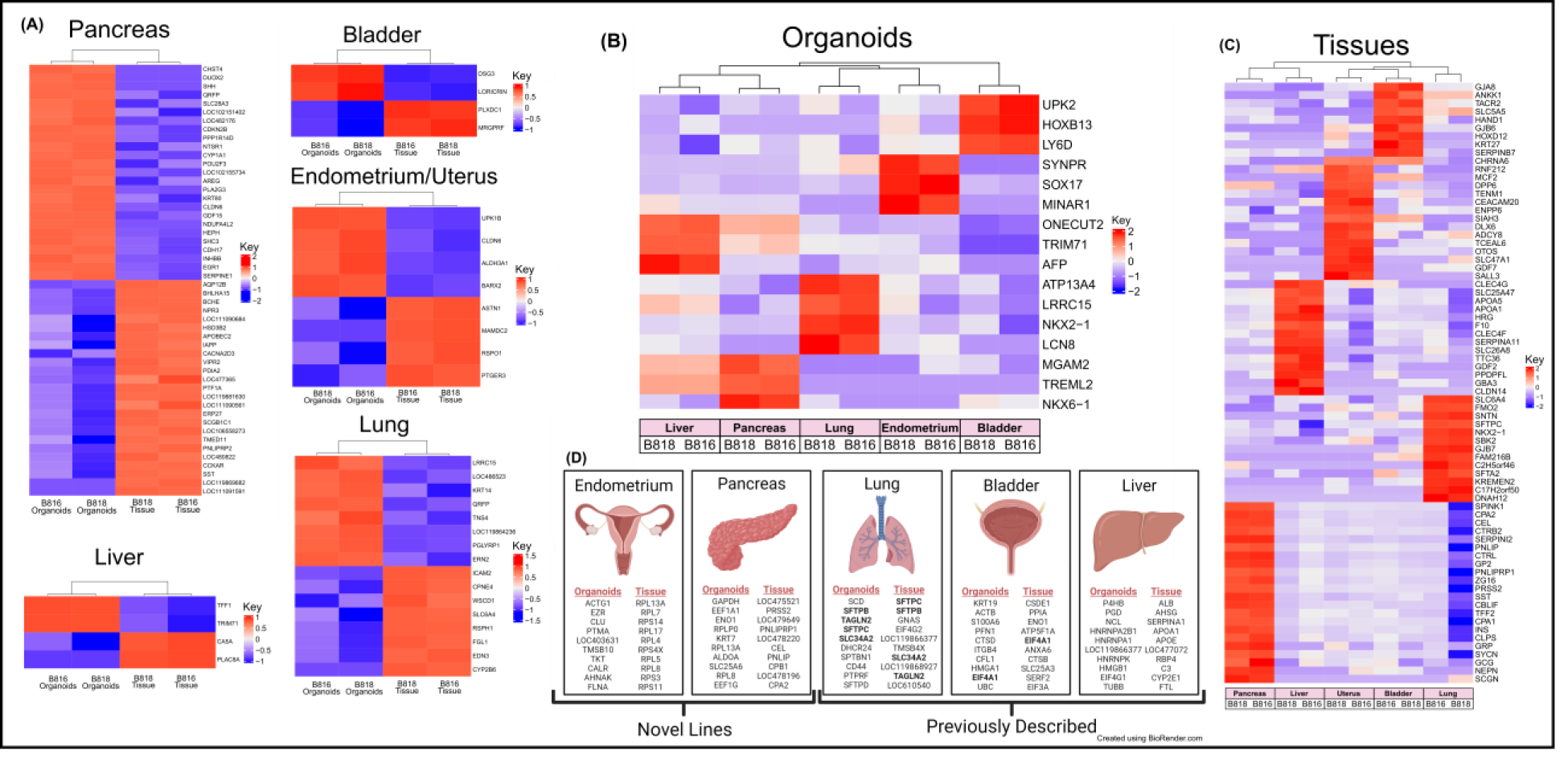
Expression of RNA and identification of tissue-specific markers for both organoids and tissues. (A) RNA heatmaps of the differentially expressed (DE) genes (FWER < 0.05) between tissues and organoids of the same organs. Tissue-specific markers were identified across the five tissues for both (B) organoids (FWER < 0.05) and (C) tissues (FWER < 0.05). Upregulated expression is red, white is neutral, and blue represents suppressed expression. (D) The ten most highly expressed tissue-specific genes from two genetically related donors for both organoids and tissues, as well as genes in common between organoids and tissues are denoted in **bold**.

**Figure 4.**
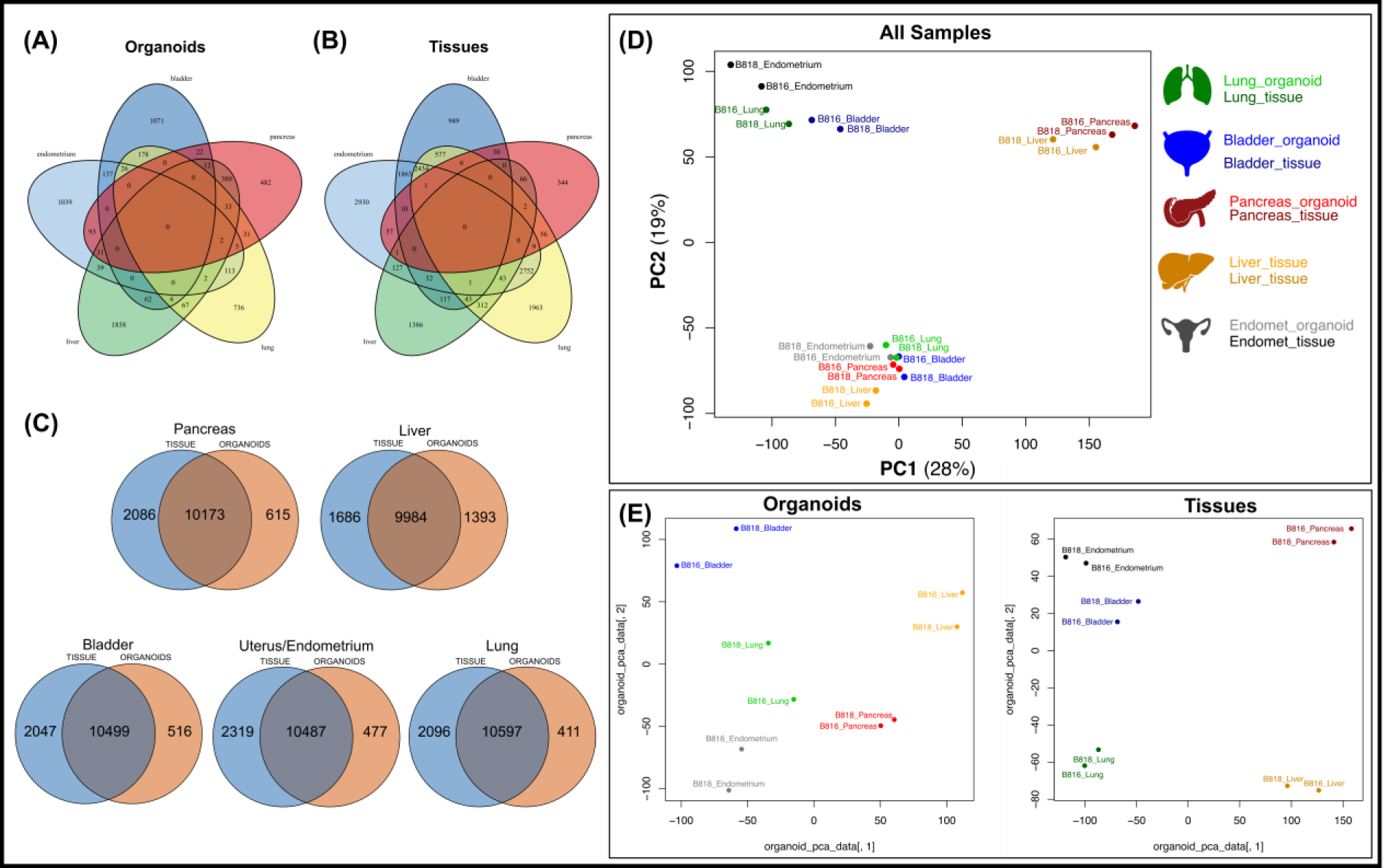
Comparison of mRNA expression similarity between organoids and tissue samples for each organ. Venn diagrams of genes expressed from both donors (B816 and B818) comparing (A) organoids and (B) tissues from the same organs. (C) Venn diagrams showing the comparison of mRNA expression between organoids and tissues from each organ. (D) Principal component analysis (PCA) plots of mRNA expression across all organoid and tissue samples. (E) PCA plots for either organoid or tissue samples. Tissue types are color coded in the legend.

**Figure 5.**
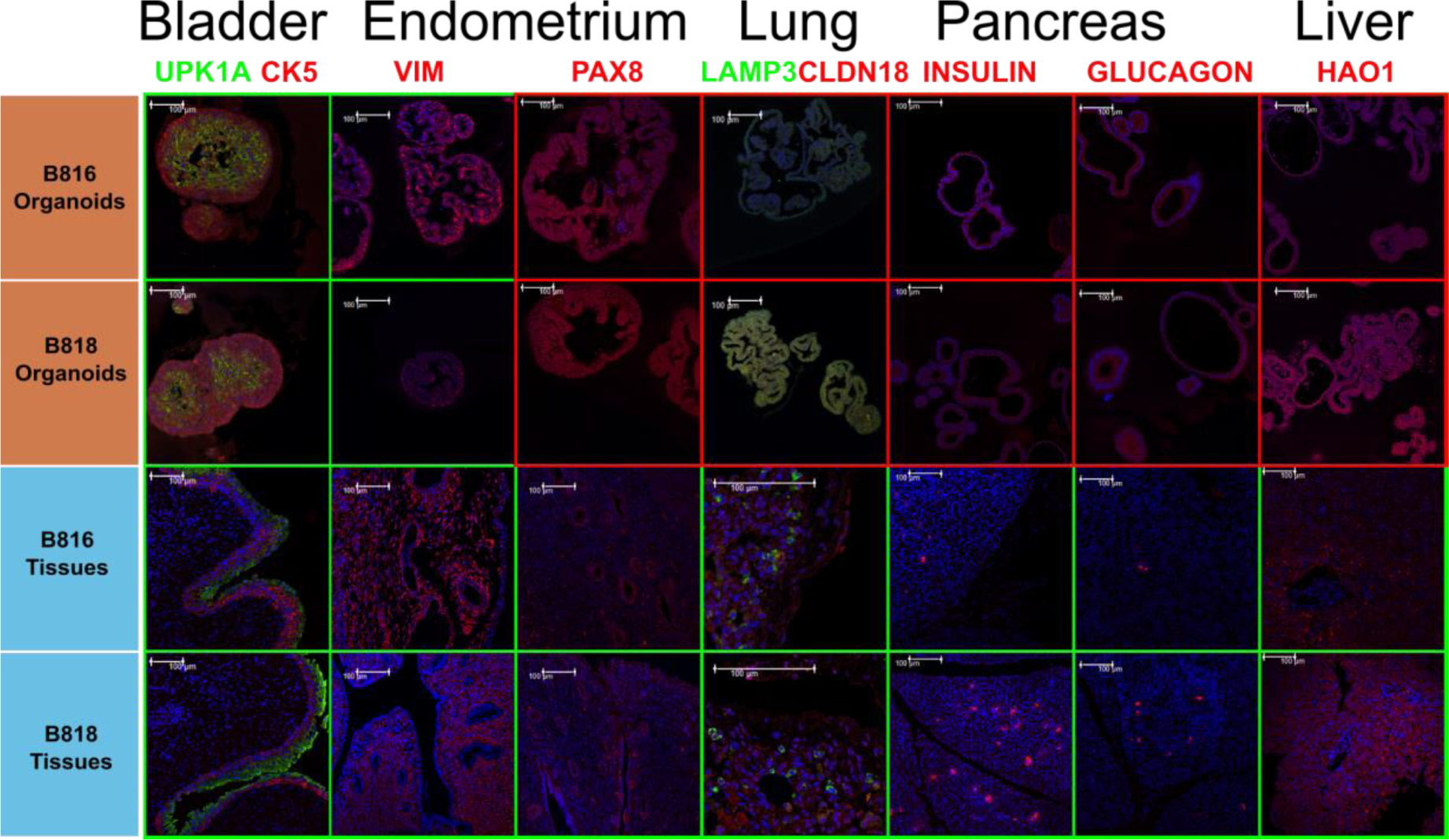
Protein characterization across tissues and organoids. Immunofluorescent staining of organoids and tissues for both donors, B816 and B818. Antibodies are labeled in the channel they appear in, and images are merged with DAPI (blue). Images were taken at either 20X or 60X magnification, scale bars are 100 μm. A green background represents positive expression of the marker while a red background indicates the lack of expression in the current field. Immunofluorescent details are listed in **Supplemental file 7**, and negative control images taken under the same conditions are shown in **Figure 5-figure supplement 1**.

#### Pancreas

The pancreatic organoids displayed two phenotypes, one resembling spheroids and another of a flowering organoid (**Figure 1**). H&E staining suggested that the culture mainly consisted of cells derived from intercalated ducts, with a few cells potentially differentiating into endocrine cells (**Figure 1**). Genes that were found to be upregulated in organoids compared to tissue included Dual oxidase 2 (DUOX2), Pyroglutamylated Rfamide peptide (QRFP), Cadherin 17 (CDH17), and Early growth response 1 (EGR1) (**Figure 3A**). Noteworthy is that Somatostatin (SST) expression was reduced in organoids suggesting our culture does not contain a significant number of neuroendocrine delta cells (**Figure 3A**). However, maltase-glucoamylase 2 (MGAM2) and NK6 homeobox 1 (NKX6-1) were also upregulated in organoids (**Figure 3B**). NKX6-1 serves as a marker of multipotent pancreatic progenitors, indicating their ability to differentiate into ductal, acinar, and endocrine cells (Wiedenmann et al., 2021). Upregulated genes in pancreas tissues included cells positive for insulin (INS), glucagon (GCG), and multiple markers characteristic of pancreatic acinar cells (**Figure 3C**). One of the most highly expressed pancreas-specific genes in the organoids included cytokeratin 7 (KRT7), indicating that most of the cells in the organoids are of epithelial origin and likely represent pancreatic ductal cells (Wiedenmann et al., 2021). Regarding pancreas-specific genes, intra-organoid comparison revealed 482 unique genes (**Figure 4A**), intra-tissue comparison revealed 344 unique genes (**Figure 4B**), and 10,173 genes were expressed in both organoids and tissues (**Figure 4C**). IF staining on pancreatic tissues identified insulin and glucagon in the islets (**Figure 5**); however, the organoids did not express measurable insulin nor glucagon.

#### Lung

The lung organoids displayed three distinct phenotypes with flowering differentiated organoids and bulbous organoids constituting most of the culture, while a small proportion had a morphology resembling alveolar structures (**Figure 1**). H&E staining suggests our culture consists of alveolar type-2 cells (AT2) and bronchial epithelial cells (**Figure 1**). The lung marker, NK2 homeobox 1 (NKX2-1) (Dost et al., 2020), was upregulated in organoids (**Figure 3B**), while Surfactant Protein B (SFTPB) and Surfactant Protein C (SFTPC) gene expression were detected and specific to both lung tissues and organoids (**Figure 3D**). Upregulated genes in organoids compared to tissues included Pyroglutamylated RFamide peptide (QRFP) and Peptidoglycan recognition protein 1 (PGLYRP1). For lung-specific genes, intra-organoid comparison revealed 736 unique genes (**Figure 4A**), intra-tissue comparisons revealed 1,963 unique genes (**Figure 4B**), and 10,597 genes were expressed in both organoids and tissues (**Figure 4C**). IF staining clearly identified alveolar type-1 cells (AT1) for Claudin 18 (CLDN18) while Lysosome-associated membrane glycoprotein 3 (LAMP3) identified AT2 cells in the tissue sections, however, organoids did not show IF signal for either (**Figure 5**).

#### Urinary Bladder

Bladder organoids displayed a single phenotype constituting round organoids without a visible lumen or internal chamber typical of spheroids of other tissues such as intestine (**Figure 1**) (Chandra et al., 2019; Gabriel et al., 2022a). H&E staining clearly showed the organoids present morphological features consistent with transitional epithelium, consisting of characteristic basal and umbrella cell layers (**Figure 1 and Figure 5**). Desmoglein 3 (DSG3), which is a basal cell marker (Elbadawy et al., 2019), and Loricrin cornified envelope precursor protein (LORICRIN), an intermediate cell marker (Lin et al., 2013), were upregulated in organoids compared to tissues (**Figure 3A**). Uroplakin (UPK) proteins are specific to terminally differentiated urothelial cells (Lu et al., 2022), and Uroplakin 2 (UPK2) were both upregulated in organoids (**Figure 3B**). The bladder-specific marker EIF4A1 was expressed in both organoids and tissues. Regarding bladder-specific genes, intra-organoid comparisons revealed 1,071 unique genes (**Figure 4A**), intra-tissue comparisons revealed 949 unique genes (**Figure 4B**), and 10,499 genes were expressed in both organoids and tissues (**Figure 4C**). The presence of umbrella cells and basal cells was confirmed by positive IF staining with Uroplakin Ia (UPK1A) and Cytokeratin 5 (CK5) (**Figure 5**) in tissues (Elbadawy et al., 2022; Kim et al., 2020a). UPK1A expression was present in the “pores” of umbrella cells, while expression of CK5 was positive and limited to the outside of the bladder organoids, which is characteristic of inverted epithelial growth in organoid cultures (**Figure 5**).

#### Liver

Liver organoids morphologically resembled the pancreatic organoids, with one phenotype resembling spheroids and another of a more flowering organoid. Cellular morphology observed under H&E evaluation suggests that most of the cells were differentiated cholangiocytes (**Figure 1**). Trefoil factor 1 (TFF1) and Tripartite Motif Containing 71 (TRIM71) were upregulated in liver organoids compared to liver tissue (**Figure 3A**). TFF1 encodes a protein critical in the regeneration of the liver after injury by promoting biliary lineage differentiation and inhibiting hepatic lineage (Hayashi et al., 2018). Single-cell RNA sequencing of the human liver described a transcriptional profile of a cell population within cholangiocytes where the DE genes included TFF1 (MacParland et al., 2018). TRIM71 was also upregulated in organoids (**Figure 3B**) and has previously been hypothesized to be involved in promoting rapid self-renewal in undifferentiated mouse embryonic stem cells (Chang et al., 2012). Liver-specific organoid markers included multiple heterogeneous nuclear ribonucleoproteins (**Figure 3D**). Albumin (ALB) was the most highly expressed liver-specific gene in tissue due to the large percentage of hepatocytes (**Figure 3D**). Regarding liver-specific genes, intra-organoid comparisons revealed 1,858 unique genes (**Figure 4A**), tissues had 1,386 unique genes (**Figure 4B**), and 9,984 genes were expressed in both organoids and tissues (**Figure 4C**). Using IF, hepatocytes in the tissue stained positively for Hydroxyacid Oxidase 1 (HAO1) (**Figure 5**) (Kampf et al., 2014). As expected, tissues showed high expression of HAO1 as hepatocytes constitute the major cell type in liver tissue while organoids were negative for this marker, suggesting they almost entirely consist of differentiated cholangiocytes, consistent with previous descriptions in other liver-derived organoid cultures (Aktas et al., 2022; Zdyrski et al., 2024).

### Insights into organ-specific genes

The usage of multiple methods (**Figure 6A**) including RNA-seq, allowed for identification of differentially expressed genes between tissues and organoids (**Figure 3A**), and assisted in the determination of the major cell populations present and absent in the organoid cell lines. We acknowledge that the organoids are exclusively composed of epithelial cells and lack other populations present in intact tissue, such as immune cells and endothelial cells. Genes expressed were identified for each tissue type (**Figure 4C**) to emphasize the similarity of expression patterns of the organoid models compared with their tissue of origin. A comparison of mRNA expression across tissues and organoids can be seen in **Figure 4**. Between 76% and 80% (**Figure 4C**) of all expressed genes overlapped for each organ between tissues and organoids.

**Figure 6.**
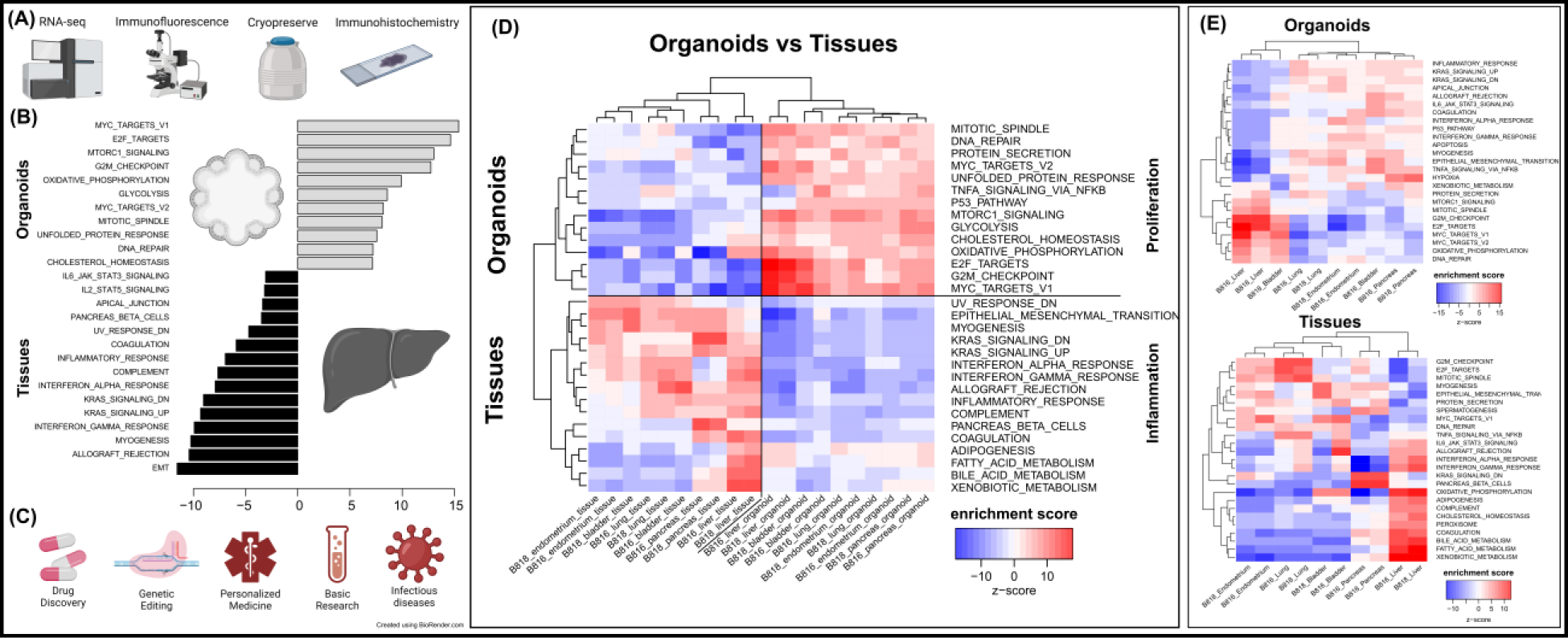
Characterization of organoids, pathway enrichment analysis, and potential biomedical applications. (A) Initial characterization and preservation methods used for the intra-donor derived canine organoid cell lines. (B) A list of the top pathways when comparing organoids to tissues. (C) A list of potential biomedical applications for canine organoid models. (D) Analysis of proliferation and inflammation pathways shown across all organoids and tissues. (E) Enriched pathways for either organoids or tissues (red = upregulated, blue = downregulated).

Principal component analysis (PCA) (**Figure 4D**) was used to visualize major sources of genetic variance for the different samples, where principal components 1 and 2 (PC1, PC2) effectively separated epithelial organoids and their tissues of origin. Furthermore, PCA across all organoids and all tissues, separately, clearly demonstrated the strongest sample clustering between genetically related animals (**Figure 4E**). Intra-organoid and intra-tissue comparisons identified upregulated genes (**Figure 3B and 3C**) and the ten most highly expressed tissue-specific genes (**Figure 3D**). The expression of unique and overlapping genes was further compared for each sample type (**Figure 4A and 4B**). Gene set enrichment analysis was used to characterize the global transcriptional programs that best characterized both organoids and tissues samples (**Figure 6E**). Stouffer integration of transcriptional pathway z-scores yielded a consensus scoring of the major global differences between paired organoids and source tissues (**Figure 6B, Supplemental file 8)**. Overall, tissue and organoid samples were best separated by transcriptional hallmarks associated with inflammation and proliferation, respectively (**Figure 6D**). This preliminary characterization will assist in determining potential applications for these novel canine organoid models for research applications (**Figure 6C**).

## DISCUSSION

### Canine organoids as biomedical models

Canines can serve as a superior model to mice for translational research applications, especially due to their tendency to develop analogous chronic diseases to those of humans and their shared similarity in lifestyle. However, using canines for translational research presents some obstacles. Their use in research can be ethically questionable and resource-intensive. Organoids can overcome some of these challenges and could potentially represent an excellent alternative to expanding the biomedical applications of the canine model. Developing novel canine organoid models will accelerate research efforts toward advanced veterinary therapeutics as well as for preclinical drug screening in human medicine. Furthermore, advances in organoid technology are being made in areas including personalized drug testing using patient-derived organoid cultures (Huch and Koo, 2015; Mullenders et al., 2019). Typically, studies report the cultivation of one tissue-specific organoid cell line, while others combine organoid lines from multiple individuals to make conclusions. By combining unrelated donors’ information, donor-to-donor variability can be neglected, thus ignoring relevant differences in patient populations. We aim to broaden the applications of the dog model in biomedicine while minimizing animal usage by developing new canine organoid lines and studying gene expression profiles across different epithelial tissues. While our tissue samples consist of various cell types (such as epithelial cells, vascular cells, and immune cells), our organoids are epithelial in origin. Comparisons between the original tissues and the derived organoids expose the constraints of current epithelial organoid models. Recently, protocols for co-culturing immune cells with organoids have become more attainable, thus enhancing the complexity and applicability of organoid models. (Chakrabarti et al., 2021).

Many laboratories utilize tissue-specific protocols and media supplemented with various growth factors to expand their organoids which can be costly and laborious (Kaushik et al., 2018; Tortorella et al., 2022). In the current study, all described cultures were grown under the same conditions using the same organoid expansion media (*Complete media with growth factors with ROCK inhibitor and GSK3β inhibitor – CMGF+ R/G*) and differentiation media (*Complete media with growth factors – CMGF+*). The use of the same media composition lends itself to future applications of co-culture or use in assembloid models where multiple organoid lines are combined and continued growth in a shared media composition is ideal. While our media composition allows for the growth and expansion of organoids and a variety of cell types, the authors acknowledge that for differentiation of certain epithelial cell types, such as within the pancreas, specific growth factors and culture conditions may need to be optimized per tissue. Additionally, many protocols emphasize tissue removal to create a suspension of stem cells during organoid isolations. Here, we utilize mechanical dissociation during isolation for the inclusion of small tissue pieces during the initial growth that we believe can benefit canine organoid expansion, assisting the most in liver cultures. The tissue is then dissociated enzymatically during organoid passages and hence removed from the culture prior to any analysis. This phenomenon we observed may be due to intercellular signaling from the stem cells still attached to damaged tissue, resulting in the release of damage-associated signals, increasing the initial growth of the stem cells, or potentially due to the availability of the extracellular matrix from the tissue at the beginning of the culture.

We report the cultivation, characterization, and comparison of five organoid lines (endometrium, pancreas, lung, bladder, and liver), two of them novel, derived from the same animal, from two genetically related canine donors of the same litter, sex, and age. Additionally, the isolation, cultivation, and media composition were identical for the five organoid cell lines, eliminating the need for tissue-specific growth factors. To the best of the authors’ knowledge, the data in this study, therefore, constitutes the most comprehensive comparison of tissue-specific expression across canine organoids available to date. These newly available canine organoids could be applied for more rapid translational applications, such as the identification of new therapeutics, the study of genetic editing technologies, and the development of better disease models. Due to the nature of the samples being derived from the same donors, these lines have the potential to be used in downstream experiments including organ-on-a-chip (Dongeun et al., 2010) and assembloid cultures (Birey et al., 2017).

### Organoid characterization and biomedical applications

#### Uterus

Previously, it has been shown that UPK1B, which was upregulated in our canine endometrial organoids, was upregulated after an endometrial biopsy, and the protein was found in glandular-epithelial cells (Kalma et al., 2009). Human endometrium organoids typically resemble a cystic-shaped organoid unlike the canine endometrial organoids, which contained large tubular structures (Boretto et al., 2017; Turco et al., 2017). One study administered hormones to the culture and noticed columnar epithelial morphology with the formation of larger vacuoles (Cindrova-Davies et al., 2021). Previously, cultures of 3D uterine glands explants and stromal cells had limited viability surviving only for four days with the resemblance of spheroids beginning to form (Stadler et al., 2009). Furthermore, 3D organotypic canine endometrium cultures have been previously described (Bartel et al., 2013), however this study simply isolated differentiated endometrial glands and stromal cells from tissue and co-cultured them for 48 hours, not attempting to proliferate or expand the cells. Endometrium organoids may also be useful in the future for investigating diseases such as endometriosis and endometrial cancers (Turco et al., 2017). Additionally, they may be used in studies related to embryo implantation into the endometrium (Rawlings et al., 2021). Finally, co-culturing endometrium organoids on three-dimensional scaffolds may provide insight into implantation studies (Cindrova-Davies et al., 2021).

#### Pancreas

The characterization of our canine pancreas-derived organoids suggests that they primarily contain intercalated pancreatic ductular cells. In addition, we believe these stem cells should have the capacity to differentiate into neuroendocrine cells but optimization of growth factors in the media may be required. In dogs, pancreatitis is by far the most common disease of the exocrine pancreas (Lim et al., 2014; Xenoulis, 2015), therefore, a healthy pancreatic canine organoid model could assist in studying the pathophysiology of pancreatitis in this species. Murine *in vivo* models have long been known to have limited translatability for modeling pancreatic cancer in humans (Bailey and Carlson, 2019). Canine pancreatic ductal organoids could potentially be used for disease modeling of pancreatic ductal adenocarcinoma (PDAC), which is one of the most lethal types of cancer in humans (Wiedenmann et al., 2021). Furthermore, differentiation of our canine pancreatic cultures can be attempted in the future since recently described methods successfully differentiated 2D canine pancreatic ductal cells into insulin- and glucagon-producing beta-like cells (Gao et al., 2022). Such applications could then be used for pancreatic hormone production and studying drug target screening and toxicological effects on the endocrine pancreas.

#### Lung

Canine lung organoids have previously been described (Shiota (Sato) et al., 2023); however, the organoid cultures we describe here contained a variety of distinct morphologies in addition to what has previously been reported. Due to the differences in the abundance of various morphologies, it was difficult to determine the expression of the least populated phenotypes using bulk RNA-sequencing. Nonetheless, phenotypic and genotypic characterization suggest that our culture may contain alveolar type 2 (AT2) cells. AT2 cells are responsible for expression of surfactant proteins in the lungs and differentiation into AT1 cells which cover more than 95% of the alveolar surface area and are crucial for gas-exchange (Barkauskas et al., 2013; Wang et al., 2018b). Lung organoid models have recently helped uncover cell pathways critical during lung repair and regeneration and to identify damage-associated transient progenitors (DATPs) which represent a distinct population of AT2-lineage cells (Choi et al., 2020). Having both bronchial epithelial cells and AT2 cell types present in our lung organoids increases the number of future potential applications of the organoids. It has been shown that the lung arises from cells expressing the NKX2-1 transcription factor (Kaushik et al., 2018), which is upregulated in our lung organoids. For example, lung organoids have previously been used in the Transwell system for studying viral uptake into cells (Zhou et al., 2018). The use of a lung organoid model derived from human pluripotent stem cells showed that AT2-like cells are susceptible to SARS-CoV-2 infection, and infection of organoids resulted in the upregulation of chemokines similar to that reported in patients with COVID-19 (Han et al., 2021). Similarly, canine lung organoids could be used for in-depth pathophysiology studies of viruses causing canine infectious respiratory disease (CIRD) complex, including canine parainfluenza virus (CPIV), canine adenovirus (CAV) type 2, and canine herpesvirus (Reagan and Sykes, 2020).

#### Urinary Bladder

Canine bladder cancer organoids were previously described and exposed to anticancer drugs to describe their potential role in research and precision medicine (Elbadawy et al., 2019). Since canine bladder cancer is a well-established model for human muscle-invasive bladder cancer (Knapp et al., 2020), canine bladder cancer organoids represent a valuable model for translational preclinical research (Minkler et al., 2021). In addition, Elbadawy et al. has recently described healthy canine bladder organoids (Elbadawy et al., 2022). This report expands the knowledge and accessibility of healthy canine bladder organoids, with those described here displaying a similar morphological phenotype to those previously described (Elbadawy et al., 2022). Using 3D patient-derived tumor organoids to predict the response to chemotherapeutic protocols has great potential in oncological precision medicine. Therefore, there is a need for healthy canine bladder organoids to serve as controls when attempting to identify novel therapeutic strategies (Yu et al., 2021).

#### Liver

Canine liver-derived organoids have been previously described from both normal and *COMMD1*-deficient dogs and were cultured to model copper storage disease, which is also known as Wilson’s disease in humans (Nantasanti et al., 2015). Our group has recently further standardized the protocol for canine hepatic organoid culture (Gabriel et al., 2022a). Based on the characterization outlined in that publication, our canine liver organoid culture is mainly comprised of differentiated cholangiocytes. The previous study describing canine hepatic organoids in expansion media showed that the organoids minimally expressed the mature hepatocyte (CYP3A12) marker while stably expressing the following markers: stem cell (CD133 and LGR5), cholangiocyte (KRT19 and SOX9), and early hepatocyte (FOXA1 and HNF4α) (Nantasanti et al., 2015). Future efforts could involve the development of media compositions, including growth factors that will enhance the differentiation of hepatic stem cells into mature, differentiated hepatocytes rather than cholangiocytes, which were first described in murine liver organoids (Huch et al., 2013), then first attempted in dogs (Nantasanti et al., 2015), further refined in dogs (Kruitwagen et al., 2020). Our group has recently investigated the ability of canine liver-derived organoid cultures to differentiate into mature hepatocytes by comparing six different media compositions (Gabriel et al., 2024). Optimization of such differentiation media could open avenues to explore their usefulness for hepatic toxicity assays in drug research, in addition to modeling various analogous cholangiopathies and hepatocellular diseases in canines.

## CONCLUSION

Applications of organoid technologies are rapidly expanding and now encompass protocols to develop reliable *in vitro* models of various diseases. Further differentiation or enrichment of certain cell populations within the organoids characterized here is warranted to expand the current scope of applications for canine organoids. We report the successful isolation, culture, and characterization of two novel canine organoid lines. These novel organoid lines will enhance future use of the technology in fields including drug development, clinical applications, and personalized medicine applications. Furthermore, a multi-tissue comparison of five canine organoid lines derived from two genetically related individuals allowed for direct evaluation of inter-organ and inter-individual variance in both in vivo and in vitro gene expression. The antibodies optimized in this study for the canine model can be used in differentiation experiments attempting to enrich specific cell types such as differentiated hepatocytes and pancreatic beta cells. Future directions, including derivation of organoids from adult and diseased canines and further characterization utilizing single-cell RNA sequencing, will help identify unique subpopulations of cells within the organoids, further increasing the applicability of these new translational *in vitro* models.

## MATERIALS AND METHODS

### Tissue collection

Dogs were used under permit (ref. IACUC-18-065), and proper Institutional Animal Care and Use Committee (IACUC) protocols were followed. For this study, two 4-week-old intact female canines were euthanized via intravenous sodium pentobarbital overdose due to unrelated reasons, and tissues were quickly harvested (donor details in **Supplemental file 3**).

Approximately 2 cm x 2 cm tissue biopsies were obtained and then rinsed three times in 10 mL of 1X Complete Chelating Solution (1X CCS, composition and further details can be found in Gabriel et al. 2022) and transferred to 6 mL of Dulbecco’s Modified Eagle Medium/Nutrient Mixture F-12 (Advanced DMEM/F12) with the addition of Pen Strep (Gabriel et al., 2022a).

### Organoid isolation and cultivation

Organoid isolation and maintenance were based on a modified protocol described by Saxena et al. in 2016, which was optimized to include the standardized culture, expansion, and harvesting of canine intestinal and hepatic organoids in Gabriel et al. 2022 (Gabriel et al., 2022a; Saxena et al., 2016). A subset of the tissue pieces was minced with a scalpel until a consistency was achieved that would fit into a 10 mL pipette, at which point the samples underwent the typical canine organoid isolation protocol. Samples were washed with 5 mL of 1X complete chelating solution (CCS) then vortexed. After the tissue settled, supernatant was removed down to the 5 mL mark and a total of five washes were done. During the last two washes, supernatant was removed down to the 3 mL mark. Next, 3 mL of 1X CCS containing the tissue sample was transferred to a 6 well plate, the sample tube was rinsed with an additional 3 mL of 1X CCS and transferred to the 6 well plate. Then, 150 µL of 0.5 M EDTA (Invitrogen, ref. 15575-038) was added and the plate was incubated at 4°C while rocking for 10 min. The sample was then transferred to a tube containing 5 mL of CCS and 2 mL of FBS and inverted. Next, the supernatant and ∼100-200 µL of tissue was transferred to an empty tube and centrifuged at 700 g for 5 min at 4°C. Supernatant was discarded and the sample was rinsed with 6 mL of DMEM, before again spinning and removing supernatant. Samples were then mixed in either of two Matrigel compositions Phenol red-free (Corning ref. 356231) or Phenol red (Corning ref. 356230) and plated in 30 µL drops in 24 well plates. Plates were incubated for ∼20 minutes at 37°C to solidify the Matrigel. Samples were expanded in our growth media (CMGF+ R/G), which is supplemented with Y-27632 ROCK inhibitor (Biogems, ref. 1293823) and a GSK3β inhibitor, Stemolecule CHIR99021 (Stemgent, ref. 04-0004). Total media volumes typically consisted of 500 µL on Monday and Wednesday, and 750 µL on Friday, any deviations are listed in **Supplemental file 1**. Passaging was done as previously described, through the addition of 500 µL TrypLE Express to 500 µL of DMEM and organoids which was then incubated in a 37°C heat bath for 10 minutes. Dissociation was stopped by dilution of the TrypLE Express with 6 mL of DMEM which was then centrifuged and removed. Cleaning of the organoids was used to replace Matrigel or change the density of the organoids in the culture. Before harvesting for characterization, both Y-27632 ROCK inhibitor and GSK3β inhibitor were retracted (CMGF+) for five days to discourage the culture from stem cell expansion and allow time for potential differentiation of cell lines (see **Supplemental file 2** for complete media details). Samples used for paraffin embedding were ensured to be plated in Phenol red-free Matrigel.

### Cryopreservation of Organoids

Two freezing medias were used to cryopreserve organoids which consisted of (1) 50% CMGF+ R/G, 40% FBS, and 10% DMSO as well as (2) Cryostor CS10 (BioLife Solutions; 210102). Recovery after Cryostor CS10 was more reliable and thus was favored. Prior to freezing, organoids were recovered from Matrigel and resuspended in an appropriate freezing media. After being placed in a 1 mL cryovial, samples were placed in the fridge for 10 minutes, then moved to a −80°C freezer overnight in a Mr. Frosty container (Nalgene; 5100-0001) filled with isopropanol, and finally stored in liquid nitrogen (−196°C) indefinitely.

### RNA extractions and sequencing

After isolation, expansion, and differentiation (between 17 and 31 days), organoids were pelleted and resuspended in 100 µL of Phosphate Buffered Saline (PBS) and transferred to a cryovial. The sample tube was flushed with 900 µL of RNAlater and subsequently added to the cryovial before being stored in liquid nitrogen (−196°C). Tissue biopsies were directly placed into cryovials containing 1 mL of RNAlater and stored in liquid nitrogen. Upon thawing, tissue samples were quickly rinsed in PBS to remove excess salts from the RNAlater solution and were immediately transferred to 800 µL of Trizol and homogenized with a pestle. Organoid samples were thawed and transferred to a 15 mL tube with 2 mL of PBS, then centrifuged at 1,200 g at 4°C for 5 min to pellet the organoids. RNAlater was removed, and 1 mL of Trizol was added to the organoids and homogenized via brief vortexing. After homogenizing, samples were stored at room temperature for 5 minutes and then centrifuged at 12,000 g at 4°C for 10 min to eliminate debris and polysaccharides. The supernatant was transferred to a new tube, and chloroform (0.2 mL chloroform per mL Trizol) was added. Samples were shaken vigorously for 20 sec and stored at room temperature for 2-3 minutes before being centrifuged at 10,000 g at 4°C for 18 minutes. The aqueous phase was transferred to a sterile 1.5 mL Rnase-free tube. Then an equal volume of 100% RNA-free EtOH was slowly added and mixed before being transferred to a Qiagen Rneasy column (Rneasy Mini kit) seated in a collection tube which was centrifuged for 30 seconds at 8,000 g. Flow-through was discarded, and the Qiagen Dnase treatment protocol was followed. Next, 500 µL of buffer RPE was added and centrifuged for 30 seconds at 8,000 g. Flow-through was again discarded, and 500 µL of buffer RPE was added and centrifuged for 2 minutes at 8,000 g. Flow-through was discarded, and columns were centrifuged for 1 minute at 8,000 g to remove the remaining buffer. RNA was eluted in 50 µL of Rnase-free water and allowed to sit for 2 minutes before being centrifuged for 1 minute at 8,000 g. Samples were centrifuged again at 8,000 g, immediately analyzed on a Nanodrop, and frozen at −80°C.

Prior to library preparation, RNA samples were quantified with an Agilent 2100 Bioanalyzer (Eukaryotic Total RNA Nano). Further quantification was done by GENEWIZ using a Qubit 2.0 Fluorometer (ThermoFisher Scientific) and a 4200 Tapestation (Agilent). An ERCC RNA Spike-In Mix kit (ThermoFisher Scientific cat. 4456740) was used to normalize total RNA prior to library preparation. A NEBNext Ultra II RNA Library Prep Kit for Illumina (New England Biolabs, Ipswich, MA, USA) was used for library preparation. mRNAs were initially enriched with Oligod(T) beads and then fragmented for 15 minutes at 94°C. Next, first and second-strand cDNA was synthesized, end-repaired, and adenylated at 3’ends, and universal adapters were ligated to cDNA fragments. This was followed by index addition and library enrichment by PCR with limited cycles. Libraries were validated on the Agilent TapeStation (Agilent Technologies, Palo Alto, CA, USA) and quantified using a Qubit 2.0 Fluorometer (ThermoFisher Scientific, Waltham, MA, USA) as well as by quantitative PCR (KAPA Biosystems, Wilmington, MA, USA). The libraries were multiplexed and clustered onto two flowcells and were loaded onto an Illumina HiSeq 4000 instrument. The samples were sequenced using a 2×150bp Paired-End (PE) configuration. The HiSeq Control Software (HCS) conducted image analysis and base calling. Raw sequence data (.bcl files) generated from Illumina HiSeq was converted into fastq files and de-multiplexed using Illumina bcl2fastq 2.20 software with one mismatch allowed for index sequence identification.

### RNA sequencing

The total number of reads from tissues and organoids ranged from ∼18 × 10^6^ to 27 × 10^6^ (**Supplemental file 4**). The average Phred quality score was 35 before quality control procedures (see *Processing, mapping, and quantification of RNA-seq libraries*). Comparisons were made between organoids and their native tissues, across organoids, and across tissues, for both B816 and B818 individuals.

### Processing, mapping, and quantification of RNA-seq libraries

Raw sequence files were inspected in FastQC v0.11 and MultiQC v1.7 (Kim et al., 2015; Li et al., 2009) to verify their quality. Barcodes were trimmed from reads and reads with a quality score < 20 were discarded from downstream analysis cutadapt v3.5 (Martin, 2011). The data set was de-duplicated with BBDuK v38.94 (https://sourceforge.net/projects/bbmap/) with a search k-mer size of 18bp. The resulting reads were passed to SortMeRNA v2.1 (Kopylova et al., 2012) to filter out rRNA sequences based on similarity with the SILVA v111 and Rfam v11.0 databases (Gardner et al., 2009; Quast et al., 2013). After each step, reads were inspected with FastQC and MultiQC to ensure the quality of the data. Prior to the alignment of reads to a dog genome with STAR v2.5, an index was created from ROS_Cfam_1.0 (RefSeq: GCF_014441545). In average for all samples, 90.5% of the reads mapped to unique targets within the reference genome (**Supplemental file 4**). Sequences from the ERCC spike-in controls were included in this index to quantify their abundances in the samples. The resulting BAM files were passed to Subread v1.6 to obtain gene-level counts via the featureCounts algorithm.

### Differential gene expression analysis

Gene counts mapped to ERCC spike-in controls by featureCounts were extracted. Then, we calculated library size scaling factors based solely on ERCC counts using edgeR v3.36 (Robinson et al., 2009) as implemented in R v4.1 (Team, 2013) and using the trimmed mean of M-values (TMM) method (Robinson and Oshlack, 2010) to normalize the ERCC counts. The scaling factors were used to normalize the gene counts and calculate log2-transformed counts per million (CPM) after adding a 0.5 as a constant to all the values.

Our goal was to detect differences in gene expression between extracted tissues and the corresponding organoids. Prior to differential gene expression analysis, visualization of transcriptional variance was explored with principal component analysis and multidimensional scaling. This unsupervised analysis revealed similarity between samples as expected by tissue-of-origin (endometrium, lung, pancreas, bladder, or liver) and type (extracted or organoid). No obvious outliers were detected during this exploratory analysis. The model under testing was expr=β_1_+β_2_ x organoid, with type indicating if the sample was an organoid or not. Gene-wise dispersions were estimated, and outlier effects were reduced with the estimateDisp function in edgeR (using the robust=T option). Negative binomial generalized linear models (GLM) were fitted for each gene, and statistical significance for the difference in mean expression was obtained by performing Bayes quasi-likelihood F-tests (glmQLFTest function in edgeR). Visualization of the results via heatmaps and Venn diagrams were generated via the ComplexHeatmap (Gu et al., 2016) and VennDiagram (Chen and Boutros, 2011) R packages. Genes unique to each organ are listed in **Supplemental file 5** and **Supplemental file 6**. Transcriptional Pathways for both tissues and organoids were analyzed by conducting single-sample Gene Set Enrichment Analysis (ssGSEA) using the VIPER algorithm (Alvarez et al., 2016). This analysis focused on 50 Hallmark gene sets which were obtained from the Molecular Signatures Database (MsigDB) (Liberzon et al., 2015). Scaled pathway enrichment scores were converted to z-scores and was visualized and clustered using the gplots R package. All data and analysis code has been made publicly available on a GitHub repository.

### Paraffin embedding and immunohistochemistry

After organoids were expanded, they were then allowed to grow in CMGF+ for five days, media was removed, and 500 µL of Formalin-acetic acid-alcohol (FAA, composition in Gabriel et al. 2022) was added to each well (Gabriel et al., 2022a). After 24 hours, FAA was replaced with 70% ethanol and samples were paraffin-embedded and mounted on slides at the Iowa State University Histopathology laboratory. Tissues were fixed in paraformaldehyde and paraffin-embedded according to standard histology procedures. Tissues and organoids were stained with hematoxylin and eosin (H&E).

For immunohistochemistry, samples were deparaffinized and rehydrated through a series of alcohol changes to deionized water. Endogenous peroxidase within the samples was then quenched using a hydrogen peroxide bath. Heat induced epitope retrieval was performed using either a tris-EDTA or citrate buffer. Immunohistochemistry (IHC) antibodies for Pan cytokeratin (Agilent, M0821), smooth actin (BioGenex, MU128-UC) and vimentin (Agilent, M0725) were used on both tissues and organoids. An indirect method of IHC staining was then carried out using a biotinylated secondary antibody followed by a streptavidin. The samples were then incubated with NovaRED™ (Vector, SK-4800) chromogen, counterstained with hematoxylin, and dehydrated. Light microscopy images were taken on a Leica Aperio GT 450 Scanner and analyzed with ImageScope (v12.4.3.5008) or on an Olympus BX40 light microscope.

### Immunofluorescence

For deparaffinization, slides were placed in xylene twice for ten minutes, then transferred to 100% ethanol twice for one minute with regular agitation. After the last alcohol wash, slides were laid on tissue paper for five minutes to dry. After deparaffinization, tissues and organoids underwent Heat Induced Epitope Retrieval (HIER) with either Citrate buffer (pH 6) or a Tris/EDTA buffer (pH 9) using a HybEZ II Oven at 75°C for two hours. After two hours, the tray was taken out of the oven, and the slides were allowed to cool with the lid off for 15 minutes. Once cool, the slides were rinsed in PBS twice for two minutes each, then rinsed in PBS for ten minutes. The tissues and organoids were permeabilized by incubation in 0.25% Triton in PBS twice for ten minutes each. After three PBS rinses, both tissues and organoids were blocked in Casein in PBS for one hour at room temperature. Tissues and organoids were incubated in a humidity chamber with their primary antibody overnight at 4°C at the appropriate concentration. The next day, the slides were again rinsed in PBS prior to the secondary being added at 1:1000 in PBS for one hour at room temperature, with slides being rinsed again. The slides were then incubated with DAPI (Sigma, D9542-1MG) at 1:500 in PBS for twenty minutes, and washed three times for ten minutes in PBS, then switched to distilled water. Fluoroshield (Sigma, F6182-20ML) was used to mount the slides, and after drying overnight, the slides were imaged on a Stellaris confocal microscope. Antibodies, dilutions, and antigen retrieval techniques for each tissue type can be seen in **Supplemental file 7**. Scale bars were added to immunofluorescent images using Leica LAS AF Lite (v. 2.6.0 build 7266).

## DATA AVAILABILITY

The RNA-seq raw reads generated in this study are available in the Sequence Read Archive (NCBI-SRA BioProject PRJNA847879) as well as the aligned files being available on the NIH ICDC (https://caninecommons.cancer.gov/#/study/ORGANOIDS01). The bioinformatic scripts are available on Github (https://github.com/chris-zdyrski/Novel_Canine_Organoids).

## ACKNOWLEDGEMENTS

We are grateful for startup funds received from Iowa State University as well as a generous donation from the Armbrust family through the Iowa State University Foundation. We would like to thank Dr. Jodi Smith for the use of her colony. We are also appreciative of Dr. Adrien Aertsens and Sichao Mao for their help with tissue collection. We thank Dr. Nicole Valenzuela for lending access to Iowa State University’s computing resources. We thank Akhil Vinithakumari for assistance with graphing of data, Ahona Mukherjee for assisting with immunofluorescence, and Dr. David Meyerholz for marker recommendations. We appreciate the timely processing of samples by the Iowa State University Pathology Department and the Iowa State University Histopathology Department.

---

**Figure 1-figure supplement 1.**
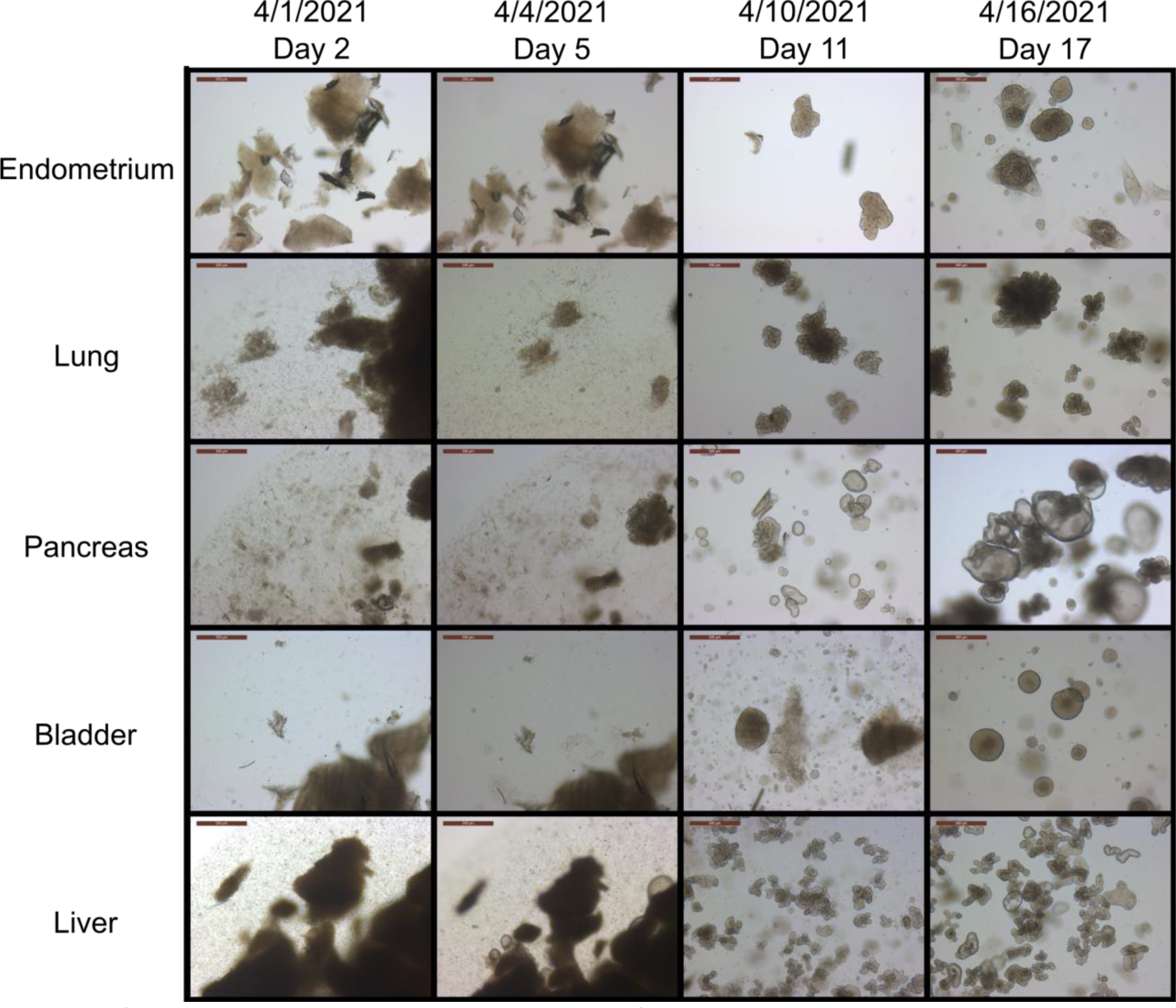
Morphogenesis of organoids over time. Light microscopy images of organoids derived from B816 tissues over time. Images spanned from two days after isolation, through a passage, to the earliest harvesting, day 17. Images were taken at 5X magnification, scale bars are 500 μm.

**Figure 2-figure supplement 1.**
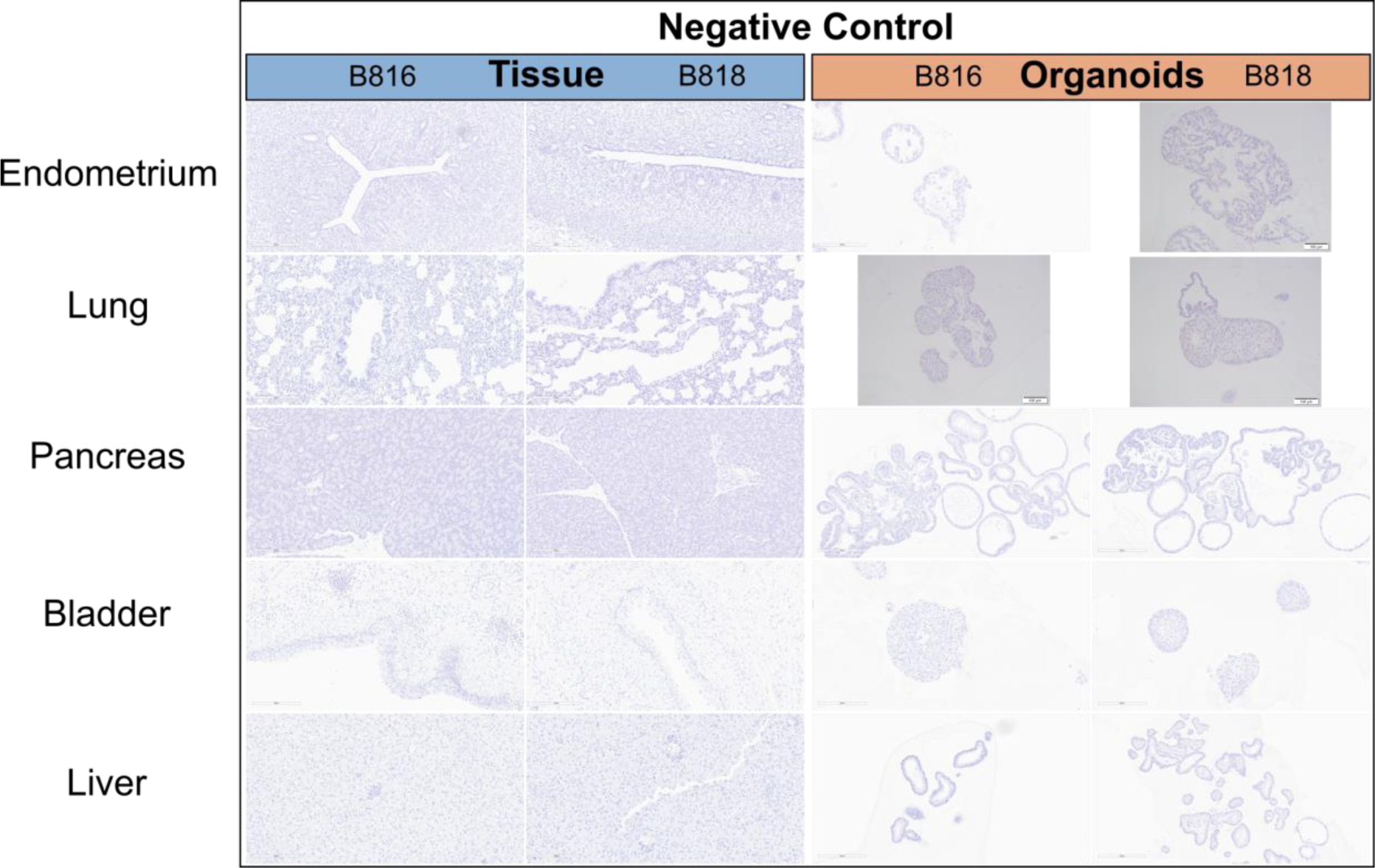
Immunohistochemistry comparison of tissues and organoids. Negative control images for immunohistochemistry staining of organoids and tissues for both donors, B816 and B818. Images were taken at 20X magnification, scale bars are in μm.

**Figure 5-figure supplement 1.**
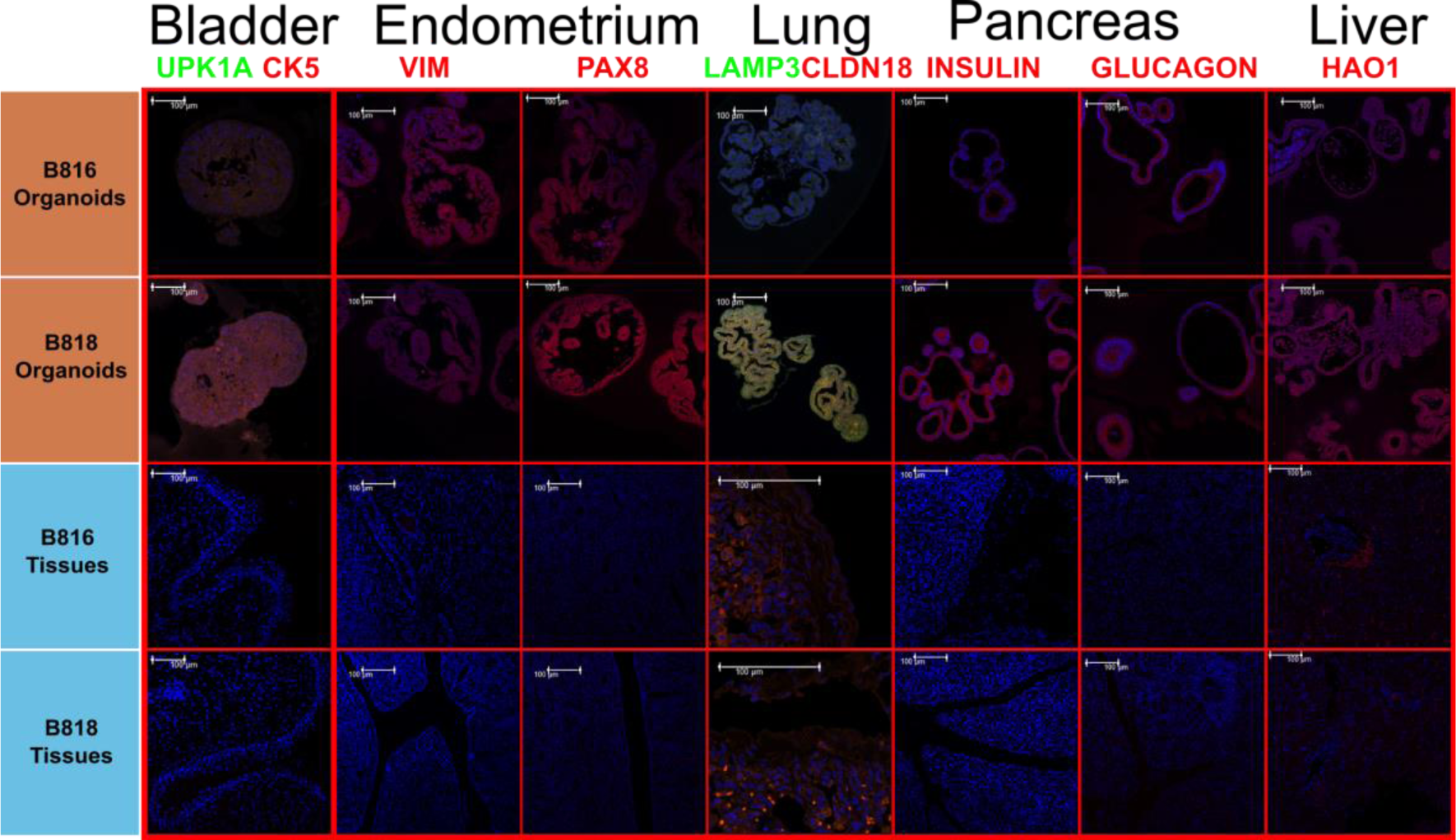
Protein characterization across tissues and organoids. Immunofluorescent staining of the negative control slides for organoids and tissues for both donors, B816 and B818. Images were merged with DAPI (blue) and were taken at either 20X or 60X magnification, scale bars are 100 μm.

**Supplemental file 1.** Growth chart for all organoid samples derived from two dogs. Descriptions include media changes, passages attempted, and total days in culture.

**Supplemental file 2.** Composition of CMGF+ culture media without the addition of RG.

**Supplemental file 3.** Donor information including age, sex, and breed from the two donors, B816 and B818.

**Supplemental file 4.** Average sequence length, total reads, quality scores, and percent of mapped reads obtained from the RNA-seq samples of organoids and tissues. Numbers are shown for the data set before and after quality control procedures (read trimming, deduplication, and filtering).

**Supplemental file 5.** The gene symbol, logCPM, and logFC values for each combination of organoids grown regarding the RNA-seq analysis. The logCPM represent the log-counts per million which are normalized expression counts. The logFC=Log fold change which compares the level of expression between two treatments.

**Supplemental file 6.** The gene symbol, logCPM, and logFC values for each combination of tissues harvested regarding the RNA-seq analysis. The logCPM represent the log-counts per million which are normalized expression counts. The logFC=Log fold change which compares the level of expression between two treatments.

**Supplemental file 7.** Antibody details, concentrations of primary and secondary antibodies, and antigen retrieval techniques used.

**Supplemental file 8.** Pathway analysis results for organoid and tissue samples from both B816 and B818. Red indicates upregulated while blue indicates downregulated pathways.

